# Acute and Chronic Neural and Glial Response to Mild Traumatic Brain Injury in the Hippocampus

**DOI:** 10.1101/2024.04.01.587620

**Authors:** Carey E. Dougan, Brandon L. Roberts, Alfred J. Crosby, Ilia Karatsoreos, Shelly R. Peyton

## Abstract

Traumatic brain injury (TBI) is an established risk factor for developing neurodegenerative disease. However, how TBI leads from acute injury to chronic neurodegeneration is limited to post-mortem models. There is a lack of connections between *in vitro* and *in vivo* TBI models that can relate injury forces to both macroscale tissue damage and brain function at the cellular level. Needle-induced cavitation (NIC) is a technique that can produce small cavitation bubbles in soft tissues, which allows us to relate small strains and strain rates in living tissue to ensuing acute and chronic cell death, tissue damage, and tissue remodeling. Here, we applied NIC to mouse brain slices to create a new model of TBI with high spatial and temporal resolution. We specifically targeted the hippocampus, which is a brain region critical for learning and memory and an area in which injury causes cognitive pathologies in humans and rodent models. By combining NIC with patch-clamp electrophysiology, we demonstrate that NIC in the Cornu Ammonis (CA)3 region of the hippocampus dynamically alters synaptic release onto CA1 pyramidal neurons in a cannabinoid 1 receptor (CB1R)-dependent manner. Further, we show that NIC induces an increase in extracellular matrix proteins associated with neural repair that is mitigated by CB1R antagonism. Together, these data lay the groundwork for advanced approaches in understanding how TBI impacts neural function at the cellular level, and the development of treatments that promote neural repair in response to brain injury.

**SIGNIFICANCE:** Current models of mild TBI (mTBI) cannot relate injury forces to both macroscale tissue damage and brain function at the cellular level. We combine a microscale injury model in *ex vivo* brain slices while simultaneously recording glutamatergic inputs onto CA1 hippocampal pyramidal neurons. Post-injury examination of chronic tissue regeneration by astrocytes allow us to connect acute neuronal signaling responses to chronic fibrosis after TBI. These studies provide a new tool for understanding the physiological and molecular responses to TBI and lay the groundwork for future experiments unraveling the synaptic mechanisms that mediate these responses seconds, minutes, and days following injury.

## INTRODUCTION

1.5 million Americans are diagnosed with traumatic brain injury (TBI) every year, and according to the Centers for Disease Control and Prevention, 5.3 million more people currently live with disabilities caused by TBI. Mild TBIs (mTBIs) account for approximately 80% of all TBI cases worldwide. Members of the military are especially at risk of cavitation-related blast associated mild traumatic brain injuries(*1–4*). When the head is exposed to subconcussive impact forces, acceleration/deceleration forces, or explosive blasts, the associated negative hydrostatic pressures can cause cavitation, or the rapid expansion of a void within the brain. There is a lack of mTBI models that relate injury forces to both macroscale tissue damage and brain function at the cellular level. Needle-induced cavitation (NIC) is a technique that induces highly localized injury to *ex vivo* brain tissue by applying fluid pressure(*5, 6*). A better understanding of cavitation damage in brain can enable better treatment options for TBI. Blast wave experiments can cause both macroscale tearing of tissue and cellular damage observed as scarring at the boundaries between white and gray matter and at blood vessel/tissue interfaces(*7, 8*). The cellular response not only leads to astrocyte-mediated glial scarring, but to accumulation of astrocyte secreted proteins during the wound healing process(*9, 10*).

Astrocytes are crucial responders to injuries throughout the central nervous system, and they have characteristic changes after injury, including astrogliosis, and changes in cell growth, size, and protein expression(*11–13*). Activated astrocytes secrete glial fibrillary acidic protein (GFAP) and extracellular matrix (ECM) proteins such as tenascin-c (TNC) and connective tissue growth factor (CTGF), which promote inflammation and neural repair(*14–19*). It is essential to identify the spatial and temporal responses of astrocyte-secreted proteins to diagnose brain injury and determine potential pathways of neurodegenerative disease progression. Although astrocyte secreted ECM proteins are known contributors to synapse formation(*20, 21*), how low strain rate cavitation acutely impacts synaptic function is unknown.

We have previously observed that NIC causes tissue damage along the hippocampus, a brain region critical for learning and memory formation(*22*). Injury to this region causes cognitive pathologies in humans and rodent models. However, the impact of NIC at the cellular level is unknown, neither acute damage to synapses nor chronic astrocyte scarring. In the present study, we combined NIC in a brain slice with patch clamp electrophysiology to investigate changes in excitatory signaling caused by the injury. We also aimed to identify potential mechanisms by which NIC impacts synaptic function, enhancing our understanding of the neural responses that follow TBI.

## MATERIALS AND METHODS

### Animals

Animal procedures and experiments were approved by the University of Massachusetts Amherst IACUC in accordance with the U.S. Public Health Service Policy and NIH Guide for the Care and Use of Laboratory Animals. Brains were collected from 4-6 week old male and female BALBc/nude, or BALBc wild type mice (Jackson Laboratories). Mice were anesthetized using isoflurane, euthanized by decapitation, and brains were removed and immediately used for NIC and organotypic brain slice preparation.

### Organotypic Brain Slicing

Immediately after brain removal and/or *ex vivo* NIC, the forebrain was blocked in a 0-4°C NMDG cutting solution (mM): NMDG 92, KCl 2.5, NaH_2_PO_4_ 1.25, NaHCO_3_ 30, sodium pyruvate 3, thiourea 2, HEPES 20, MgSO_4_ 10, CaCl_2_ 0.5, glucose 25, sucrose 20, pH 7.4 with HCl (*23*). Forebrains were mounted individually, or as two brains adjacent to each other, and sectioned simultaneously at 300 µm with a sapphire knife (Delaware Diamond Knives, Wilmington, DE) on a vibratome (Leica VS1200), yielding 3-5 hippocampal slices per mouse. Slices were then used for tissue culture or allowed to recover for patch-clamp electrophysiology experiments.

### Tissue/organotypic slice culture

Slices were placed on tissue culture inserts (EMD Millipore, Burlington, MA USA) and cultured at 37°C with 5% CO_2_ for up to 14 days with slice media(*24*).

### Brain Slice Electrophysiology

Slices were allowed to recover from slicing for 30 min at RT in recording artificial cerebrospinal fluid (aCSF) (mM): 124 NaCl, 3.7 KCl, 2.6 NaH_2_PO_4_, 26 NaHCO_3_, 2 CaCl_2_, 2 MgSO_4_, 10 glucose, and bubbled using 95% 02/5% C02. For recording, slices were transferred to a perfusion chamber containing aCSF maintained at 34-37°C. Neurons were visualized with an Olympus BX51WI microscope. Recording electrodes were back-filled with internal solutions as follows: 125 mM K-gluconate, 10 mM KCl, 10 mM NaCl, 5 mM HEPES, 10 mM EGTA, 3 mM NaATP and 0.25 mM NaGTP. Patch electrodes (3-5 MΩ) were guided to neurons with an MPC-200-ROE controller and MP285 mechanical manipulator (Sutter Instruments, Novato, CA USA). Neurons were held at V_Hold_ = -70 mV for sEPSC recordings. Recordings were collected with a UPC-10 USB dual digital amplifier and Patchmaster NEXT software. All electrophysiology reagents were purchases from Sigma Aldrich, and AM4113 was purchased from Tocris Bioscience.

### NIC on *Ex Vivo* Mouse Brain Slices

A custom pulled pipette (∼5 µm diameter tip) backfilled with aCSF was inserted ∼2-3 cell layers deep in the hippocampus and monitored with an Olympus BX51WI microscope. The NIC pipette was pressurized with the aCSF using a syringe pump (World Precision Instruments) at 5 μL/min until a bubble injury occurred at the tip of the needle observed by a drop or leveling off of pressure. Pressure was monitored in real time using a pressure sensor (Omega Engineering). Slices were then bisected along midline, and the injured hemisphere was placed on tissue culture inserts (EMD Millipore) and cultured at 37°C with 5% CO_2_ for 1-3 days in slice media(*24*).

### Slice Processing and Conditioned Media Collection

Day 0 and 3 NIC injured and sham slices on culture inserts were removed from incubator and transferred to a fresh 6 well plate and gently rinsed with 1X PBS to detach slices from insert membrane. Slices were places in 4% paraformaldehyde (Thermo Fisher) and placed in the 4°C fridge overnight. Conditioned media from injured and sham slices and syringe filtered (0.45 µm) (Fisher Scientific), flash frozen, and stored in the -80°C freezer.

### Mouse Brain Immunohistochemistry

Fixed sham and injured slices were rinsed with 1X PBS (3X) and permeabilized and blocked with 0.3% (v/v) Triton X-100/in Intercept blocking buffer (LI-COR Biosciences, 927-70001, Lincoln, NE USA) with 10% donkey serum (Abcam, ab7475) for 1 hour at 4°C. Slices were rinsed in 1X PBS (3X) and stained with primary antibodies: GFAP (Abcam, ab7260), TNC rat Invitrogen, MA1-26778, TSP (Novus, NBP1-52410, St. Charles, MO USA), and CTGF (Thermo Fisher, MA5-31420) diluted in Intercept blocking buffer at 1:250 overnight at 4°C. Slices were rinsed with 1X PBS (3X) and stained with secondary antibodies: donkey anti-rat 570 (Jackson ImmunoResearch Laboratories: 712-295-153, West Grove, PA USA), donkey anti-goat 405 (Jackson ImmunoResearch Laboratories: 705-475-747), donkey anti-rabbit 647 (Jackson ImmunoResearch Laboratories: 711-605-152), and donkey anti-mouse 488 (Jackson ImmunoResearch Laboratories:715-545-150) at 1:400 for 1 hour at 4°C. Slices were rinsed with 1X PBS (3X) and mounted on charged slides (Genesee Scientific, 29-107, Rochester, NY USA) with Gelvatol and covered with 24 x 60 mm coverslips (Corning, 12-553-472, Corning, NY USA) and sealed with nail polish (Sally Hansen, New York, NY USA). Slides were imaged on a Spinning Disc Observer Z1 microscope (Carl Zeiss).

### Statistical Analysis

Electrophysiology data was analyzed in Patchmaster NEXT or converted with ABF Utility (Synaptosoft, Fort Lee, NJ USA) for analysis in Clampfit (Molecular Devices) and/or MiniAnalysis (Synaptosoft). Datasets were tested for normality before statistical analysis. Comparisons of data between conditions was done using a one-way ANOVA or a Kruskal-Wallas test for datasets that failed normality. Statistics were calculated with Prism 9 (Graphpad) and Python 3.11.

## RESULTS

### Precision Model for Studying Traumatic Brain Injury

The hippocampus is highly involved in memory formation, which is often disrupted after TBI(*25, 26*). However, many previous approaches use more generalized TBI models, such as mechanical-force weight drop, fluid percussion, and blast-induced injury(*27*). Needle-induced cavitation (NIC) is a technique allowing for highly localized brain injury(*5, 28*). Here, we are the first to apply hippocampus specific NIC to an organotypic brain slice with µm resolution, while simultaneously using patch-clamp electrophysiology to continuously (sampling at µs resolution) measure synaptic responses before, during, and after injury in the hippocampus (Fig. 1a,b). For this technique, we simultaneously placed an NIC injury pipette into the CA3 region of the hippocampus and used whole-cell patch clamp to measure synaptic responses in CA1 pyramidal neurons (Fig. 1a). The pressure in the NIC pipette was continuously measured until an instability occurs at a critical pressure *P*_*c*_ (Fig. 1b-d). Immediately following injury, the tissue closes around the ruptured injury cite, which can be localized through inclusion of rhodium beads in the pipette (Fig. 1e). Needle puncture was validated through microscopic imaging of the needle insertion process.

**Figure 1.**
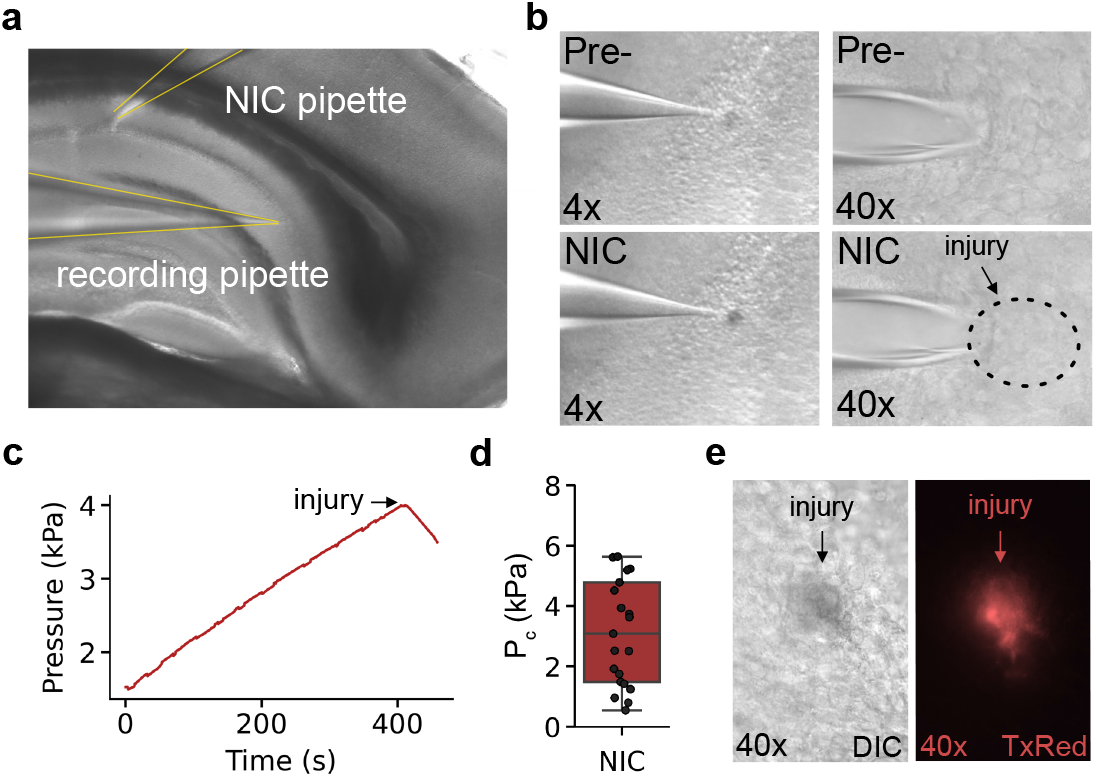
Characterization of Novel Organotypic Slice NIC Model for Studying Synaptic Function Following Brain Injury. **a** Image of simultaneous recordings of injury force and excitatory responses in CA1 hippocampal pyramidal neurons (Bottom) **b** Live imaging just prior (top) and at exact time of injury (bottom) at 4x (left) and 40x (right) magnification. **c** Representative recording trace and **d** mean critical pressure (Pc) required to evoke an NIC injury event. **e** Image of injury site after NIC pipette removal (left) labeled with rhodium beads from injury pipette (right). Boxplot represents median, inner- and outer-quartile range and are displayed with individual datapoints.

### *Ex Vivo* NIC Transiently Decreases Glutamate Release onto CA1 Pyramidal Neurons

It is unclear how synaptic function in the hippocampus is altered during and immediately following TBI. To better understand the circuit level changes in the hippocampus immediately following TBI, an approach with high temporal resolution is needed. Here we incorporated a novel approach to studying TBI (Fig. 1) and used whole-cell patch-clamp techniques to measure spontaneous excitatory post-synaptic currents (sEPSCs) onto CA1 pyramidal neurons (Fig. 2a). We found that in the 1–2-minute period following NIC injury, the frequency of excitatory events onto CA1 pyramidal neurons was greatly reduced (Fig. 2a-d). However, this transient decrease in sEPSC frequency was followed by a strong rebound in excitatory activity 5-10 minutes proceeding the initial injury (Fig. 2b-d). As measured by sEPSCs amplitude, there were no indications that the injury induced immediate postsynaptic effects on excitatory transmission (Fig. 2e). These data suggest that synaptic responses to brain injury are highly dynamic within the hippocampus and a potential protective mechanism occurs directly at the time of injury. Further, we show that these changes are due to presynaptic inputs, likely projections from the CA3 Schaffer collaterals, and are not a postsynaptic response to injury.

**Figure 2.**
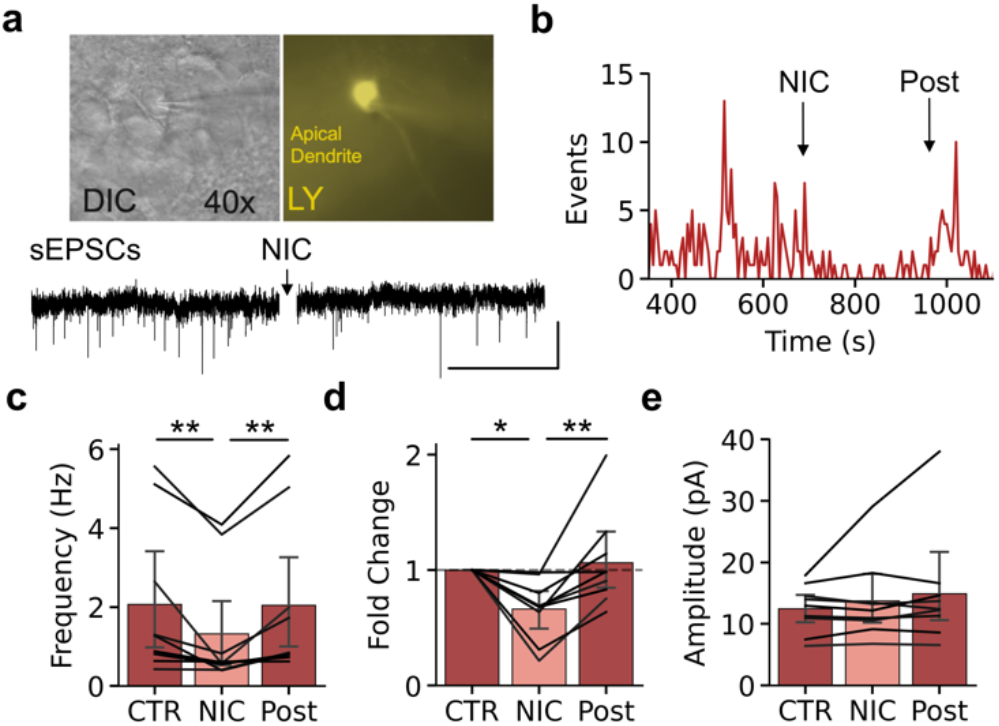
Decrease in Glutamate Release Post-NIC. **a** Representative cell (left) with lucifer yellow (right) in patch pipette for visual identification of a CA1 pyramidal neuron and a representative trace of sEPSCs before and immediately following NIC injury (bottom). **b** Histogram in 10s bins of average sEPSC events before, during, and after NIC. **c** Average sEPSC frequency (F_(2,16)_ = 8.39, p = 0.003), **d** frequency expressed as fold change (F_(2,16)_ = 7.67, p = 0.005), and **e** amplitude (F_(2,16)_ = 1.15, p = 0.34) before, immediately after, and 5-7min post NIC. Error bars represent +/- SEM. One-way repeated measures ANOVA with Tukey posthoc analysis. *p < 0.05.

### CB1 Receptor Activation Mediates Synaptic Release in Response to Brain Injury

Cannabinoid receptor 1 (CB1R) is a known mediator of excitatory synaptic release and previous work demonstrates that CB1R activation decreases excessive glutamate release in the hippocampus(*29*). We hypothesized that CB1R activation may mediate the transient decrease in glutamate release observed directly after NIC induced hippocampal injury. To test this, we repeated the experiments from Figure 2 and measured sEPSCs in the presence of CB1R antagonist AM4113. Surprisingly, we found that pretreatment with the bath application of AM4113 (100 nM) was sufficient to block the immediate decrease in excitatory release following NIC (Fig. 3a-c). There were also no apparent postsynaptic effects of excitatory signaling as measured by sEPSC amplitude (Fig. 3d). Together, these data suggest that eCB signaling is a core component in mediating synaptic function in response to brain injury.

**Figure 3.**
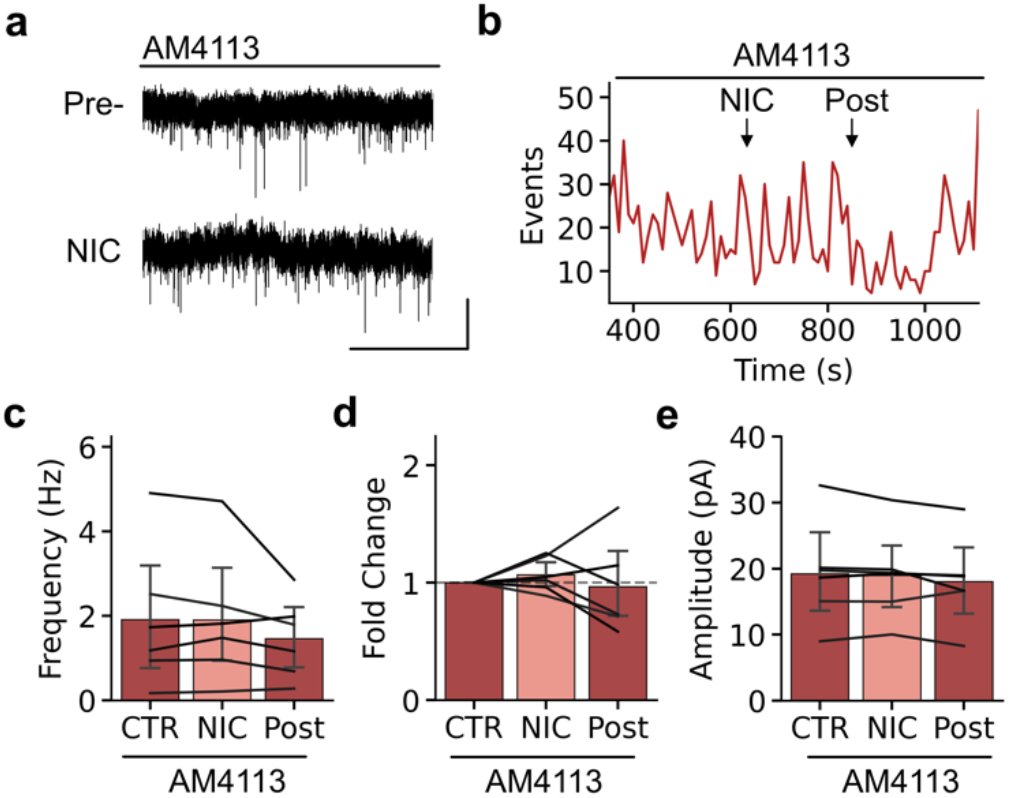
CB1R Blockade Inhibits Transient Decreases in Synaptic Transmission Following NIC Injury in the Hippocampus. **a** Representative trace of sEPSC recording in the presence of CB1 antagonist AM4113 before and immediately post-NIC injury. **b** histogram in 10s bins of average sEPSC events with bath application of CB1R antagonist AM4113 (100nM) before, during, and after NIC. **c** Average sEPSC frequency (F_(2,10)_ = 1.81, p = 0.213), **d** frequency expressed as fold change (F_(2,10)_ = 0.366, p = 0.702), and **e** amplitude (F_(2,10)_ = 1.62, p = 0.246) before, immediately after, and 5-7min post NIC. Error bars represent +/- SEM. One-way repeated measures ANOVA.

### Hippocampal Astrocyte Activation is Higher 72 Hours Post-Injury

In response to injury, astrocytes are activated to varying degrees with corresponding changes in gene expression, morphology, proliferation, and contribution to repair and remodeling(*13*). GFAP expression is an established measure of astrocyte activation. In the hippocampus injured region of mouse brain slices, astrocyte activation was higher than in control slices of brain tissue (Fig. 4a-b).

**Figure 4.**
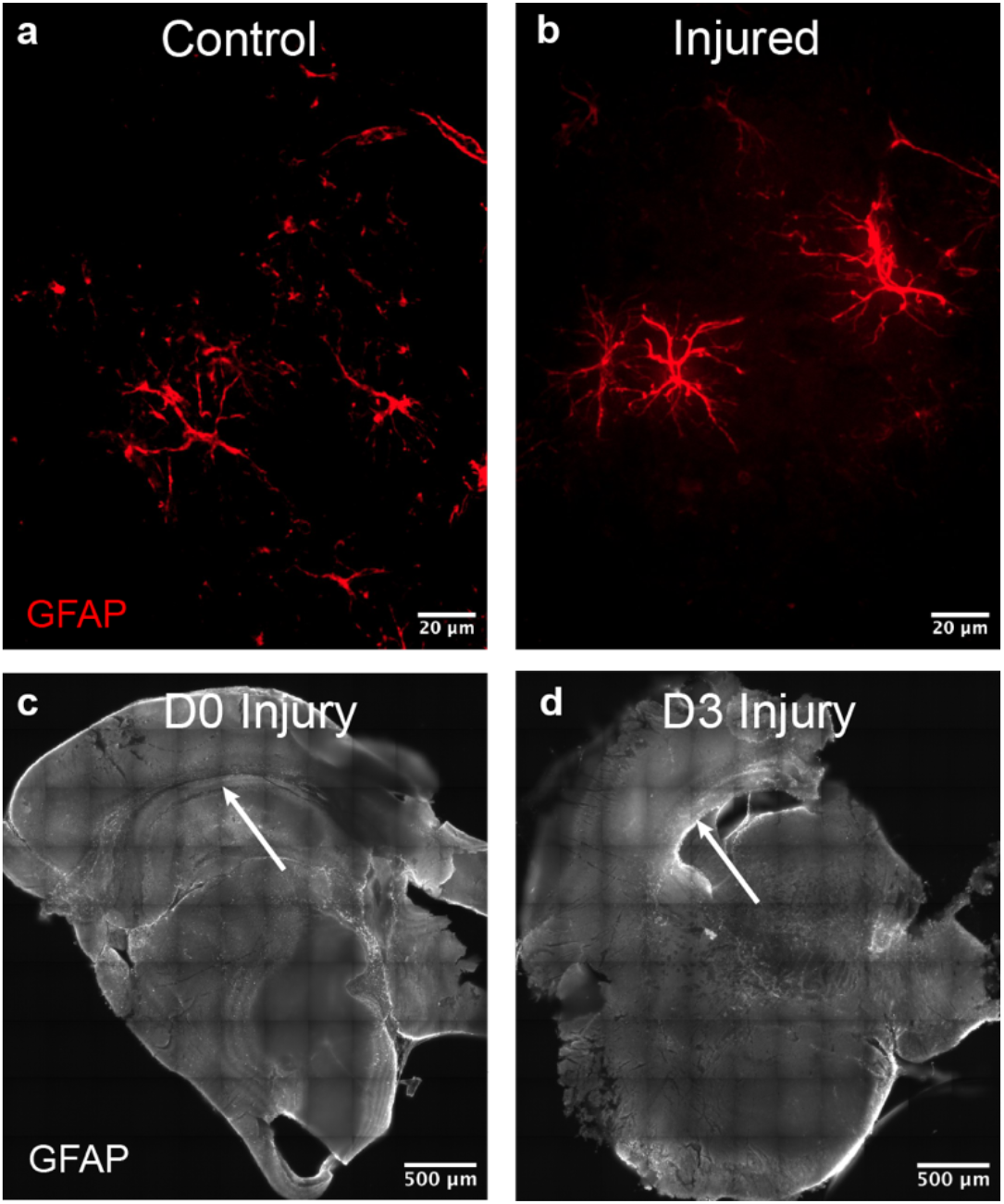
Astrocyte Activation in NIC Injured Brain Slices. Representative astrocyte activation (GFAP-red) of **a** control and **b** NIC injured brain slices at Day 0. Representative images of astrocyte activation (GFAP-white) in the hippocampus (white arrows) injured region of brain slices at **c** Day 0 (D0) and **d** Day 3 (D3) post-injury.

Astrocyte activation increased over the course of 72 hours post-injury compared to Day 0 injured slices (Fig. 4c-d). This suggests that astrocytes have an acute response to NIC injury that continues for at least 72 hours.

### Extracellular Matrix Protein Upregulation Mitigated by CB1R Antagonist

The matricellular protein families: secreted protein acidic and rich in cysteine (SPARC), tenascin-c (TNC), thrombospondin (TSP), and CCN (CYR61/CTGF/NOV), are upregulated in reactive astrocytes following injury or disease(*12*). To test this, we used immunohistochemistry to assess how several of these proteins changed following NIC. We expected post-injury acute astrocyte activation, and increased activation and secretion of extracellular proteins: GFAP, TNC, and CTGF in the days following injury. Immunohistochemistry of Day 0 slices indicated similar levels of TNC and CTGF with a slight increase in GFAP expression for NIC injured (with and without the CB1R antagonist) versus sham slices (Fig. 5). Interestingly, after 72 hours, GFAP, TNC, and CTGF were upregulated in NIC injured versus sham slices (Fig. 4c-d & Fig. 5). The addition of a CB1R antagonist at the time of injury decreased the upregulation of remodeling proteins at 72 hours post-injury (Fig. 5).

**Figure 5.**
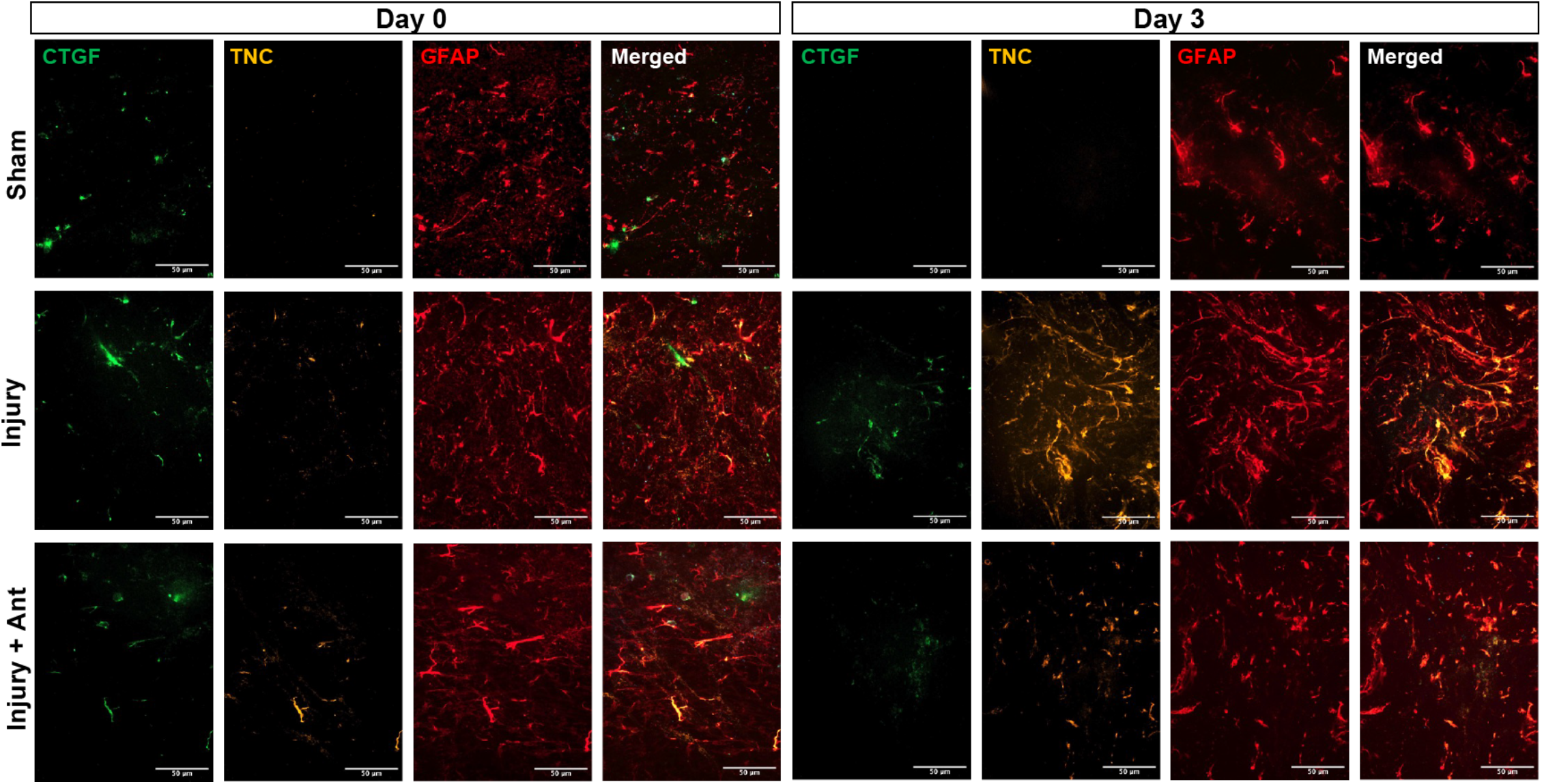
Day 0 vs. Day 3 Protein Staining. Sham, injury, and injury + antagonist IHC staining images for connective tissue growth factor (CTGF-green), tenascin-C (TNC-orange), glial fibrillary acidic protein (GFAP-red), and merged at Day 0 vs. Day 3. Scale bar is 50 µm.

## DISCUSSION

Progress in understanding the immediate cellular sequelae of TBI has been hampered by a lack of tools. In this study, we aimed to address this by combining several technical approaches that revealed important new information about the short-term effects of TBI on hippocampal circuits. *Ex vivo* cultured brain slices have been used to study neurological function, disease, and injury(*30–34*). These cultured or organotypic slices maintain the cellular complexity and many of the 3D structural properties of brain tissue, which allows a more physiologically relevant approach to understanding the longer term (more than 30 minutes) biochemical mechanisms associated with TBI. TBI is linked to increased risk for a variety of neurodegenerative diseases including dementia, Alzheimer’s disease, and chronic traumatic encephalopathy (CTE)(*35–38*). Understanding the cavitation phenomenon will provide better understanding of the causation of these diseases, and therefore may help scientists/engineers develop early detection methodologies, treatments, and prevention measures. This information will elucidate the role of CTGF and TNC in long term brain remodeling, which might be important for understanding the link between TBI and neurodegenerative diseases such as Alzheimer’s Disease and epilepsy(*39–42*). These findings lay the groundwork for numerous future studies that can utilize this new technique to identify the mechanisms underlying traumatic brain injury and neurodegenerative disease.

Although TBI is a risk factor for numerous neurological disorders, many of the underlying mechanism on how TBI impacts neural function are unknown. Current models of TBI often lack reproducibility, have high mortality rates, and/or involve surgical techniques. Further, many of these techniques lack the ability to measure changes in neural function with high temporal resolution. NIC allows precise spatial control over hippocampal injury in mouse brain slices. Understanding how TBI impacts short- and long-term astrocyte responses and synaptic function lays the groundwork for advanced approaches in understanding how TBI impacts neural function, and the development of treatments that promote TBI repair and prevent neurodegenerative disease.

The impact of cellular damage on hippocampal memory formation after TBI is often measured by changes in long-term potentiation (LTP) at the synapse post-injury (*43, 44*). However, current models of TBI lack the temporal and spatial resolution necessary for measuring synaptic function in real-time. Conflicting reports suggest that glutamate levels and signaling either increase or decrease immediately following hippocampal injury (*45–49*). Current studies rely on techniques such as microdialysis and microarrays to measure extracellular glutamate levels on the order of minutes, hours, and days post-injury, and indirectly infer how this may impact neural function. These studies have demonstrated that there is unregulated release of glutamate, and a buildup of extracellular glutamate following TBI (*50, 51*). However, other reports using magnetic resonance spectroscopy (MRS) show a decrease in glutamate in the first few hours to days following TBI (*52, 53*). This discrepancy is attributed to microdialysis measuring extracellular glutamate, while MRS measures both intra- and extracellular glutamate levels. However, neither approach addresses the functional synaptic responses immediately pre- and post-injury. Combining NIC with patch-clamp electrophysiology, we measured in real time that excitatory release is highly dynamic at the onset of injury.

Here we show small scale CA3 NIC injury clearly reduces excitatory glutamate release onto downstream CA1 pyramidal neurons for ∼1-2 minutes post-injury. We speculate that this may contribute to short-term memory loss of events leading up to TBI, such as after a concussion. This is followed by a marked increase in glutamate release several minutes after the injury event. Interestingly, excessive glutamate release in the hippocampus can lead to seizures, and up to 10% of concussion patients develop epilepsy following the injury event. Thus, NIC applied directly to an organotypic brain slice is an effective technique for measuring changes in neural function with high spatial and temporal resolution.

The central nervous system (CNS) endocannabinoid system is a network of neurotransmitters and receptors that regulates many physiological and cognitive processes. The endocannabinoid system is essential in maintaining the excitatory-inhibitory balance in the hippocampus and excessive glutamate release results in the activation of CB1Rs, which serve as a negative feedback mechanism. There are two key cannabinoid receptors in the brain: cannabinoid receptor 1 (CB1R) and cannabinoid receptor 2 (CB2R). In the brain, CB1R is primarily expressed on the presynaptic neuron and on astrocytes, while CB2R is expressed on microglia (*54, 55*). In neurons, CB1R activation leads to a decrease in presynaptic glutamate release. The accumulation of eCBs in response to injury, anti-inflammatory effects, and their role in neurogenesis, suggest that eCBs may contribute to a neuroregenerative response post-TBI(*56*). One consequence of eCB effects on astrocytes is the increase in cytosolic Ca^2+^ signals through cannabinoid (CB1) activation. In the context of TBI where increased neuronal firing and excitotoxicity are common, activation of the CB1R may mediate neurotransmitter release and provide a negative feedback mechanism in response to high levels of neural activity. Interactions between eCBs and neurons, astrocytes, and microglia promote anti-inflammatory and neuroprotective effects post-TBI (*57, 58*). We show that the transient decrease in glutamate release following NIC is due to CB1R-mediated buffering at excitatory synapses. We observe that pre-treatment with the CB1R antagonist AM4113 greatly attenuates the transient decrease in excitatory release following injury (Fig. 5). Additionally, CB1R inhibition increases basal glutamate release prior to injury. Future work is necessary to determine the exact mechanisms underlying the relationship between the eCB system and hippocampal function following injury.

eCB-mediated bidirectional communication between astrocytes and neurons has been demonstrated to significantly impact synaptic plasticity, however the role of eCBs in astrocyte remodeling of brain tissue is largely unexplored. In some cases, astroglial CB1R antagonism reduces GFAP protein expression and promotes an anti-inflammatory state by simultaneously lowering levels of pro-inflammatory cytokines(*59, 60*). eCB signaling acts on CB1Rs on astrocytes, increasing intracellular calcium and signaling back to neurons in a feedback loop to release glutamate, which might be important in remodeling (*59, 61*). Surprisingly, when brain slices were treated with a CB1R antagonist prior to NIC, there were reduced levels of astrocyte secreted remodeling proteins after 72 hours than in NIC injured slices without the antagonist. We speculate that either the antagonist interrupted the repair mechanism post-injury, or that CB1 signaling after washing away the antagonist overcompensated and interrupted the remodeling process. Future work is necessary to determine the role of endocannabinoid-mediated effects on remodeling post-TBI.

Astrocytes make up ∼30% of the brain cell population. In addition to their many functions in the healthy central nervous system, astrocytes respond to CNS damage and disease through a process called astrogliosis, or changes in the molecular and functional levels in response to pathologies. Intermediate filaments (IF) are networks of long strands of proteins that provide mechanical support for cells. GFAP is the principal astrocyte IF protein in astrocytes, along with vimentin, synemin, and lamin(*62*). The upregulation of matricellular proteins by reactive astrocytes in response to injury is dependent on the type, location, and severity of insult. Given their role in remodeling the microenvironment surrounding regions of brain injury, astrocyte-secreted extracellular proteins may represent important therapeutic targets for CNS repair. GFAP, TNC, and CTGF are upregulated in the hippocampus injured region of mouse brain slices over sham slices after 72 hours. Conversely, prolonged activation of astrocytes may result in long-term accumulation of extracellular proteins that may promote the progression of neurodegenerative diseases and/or the increased risk for delayed epileptic episodes associated with TBI (*63*). Future work is necessary to determine whether the protein upregulation is reversible or if there is a link to acute mild TBI and chronic neurodegenerative pathologies.

## CONCLUSIONS

In this research, we developed a novel technique to perform NIC in an *ex vivo* brain slice simultaneously recording glutamatergic inputs onto CA1 hippocampal pyramidal neurons. The high spatial and temporal resolution of this technique has allowed us to fill a major gap in knowledge in the understanding of how acute injury to the hippocampus alters glutamate release. There is conflicting data in the literature on whether acute injury to the hippocampus induces an increase or decrease in glutamate release. Using our newly developed technique, we established that NIC induces an immediate presynaptic reduction in glutamate release for the first 1-2 minutes followed by a long-term increase in excitatory activity (5-10 minutes post injury). This suggests that conflicting reports in the literature are correct but did not have the temporal resolution to identify this mechanism. Using a CB1R antagonist, we show that an initial decrease in excitatory activity is mediated by an eCB feedback mechanism. We posit that this may be to protect against excessive excitatory activity immediately following acute injury to the hippocampus and leads to lower levels of protein remodeling post-injury. These studies provide a new tool for understanding the physiological and molecular responses to TBI and lay the groundwork for future experiments unraveling the synaptic mechanisms that mediate these responses seconds, minutes, and days following injury.

## AUTHOR CONTRIBUTIONS

C.E.D and B.L.R. contributed to conceptual design, drafting the manuscript, data collection, analysis, and data interpretation. I.K. and S.R.P. contributed to conceptual design, data interpretation, and editing the manuscript.

## ACKNOWLEDGMENTS

This research was supported by a grant from the UMass Inspiration Award for Neuroscience & Technology to C.E.D. and B.L.R. C.E.D. was supported by an NIH funded Chemistry Biology Interface Fellowship (T32 GM008515 and T32 GM139789), and a Spaulding-Smith Fellowship from the UMass Amherst Graduate School. S.R.P. was supported by an Armstrong Professorship from UMass Amherst. I.N.K was supported by an NSF CAREER award (1553067) and an NIH R01 (DK119811).

